# Lost in Space: The Impact of Traditional Neuroimaging Methods on the Spatial Localization of Cortical Areas

**DOI:** 10.1101/255620

**Authors:** Timothy S. Coalson, David C. Van Essen, Matthew F. Glasser

## Abstract

Localizing human brain functions is a long-standing goal in systems neuroscience. Towards this goal, neuroimaging studies have traditionally used volume-based smoothing, registered data to volume-based standard spaces, and reported results relative to volume-based parcellations. A novel 360-area surface-based cortical parcellation was recently generated using multimodal data from the Human Connectome Project (HCP), and a volume-based version of this parcellation has frequently been requested for use with traditional volume-based analyses. However, given the major methodological differences between traditional volumetric and HCP-style processing, the utility and interpretability of such an altered parcellation must first be established. By starting from automatically generated individual-subject parcellations and processing them with different methodological approaches, we show that traditional processing steps, especially volume-based smoothing and registration, substantially degrade cortical area localization when compared to surface-based approaches. We also show that surface-based registration using features closely tied to cortical areas, rather than to folding patterns alone, improves the alignment of areas, and that the benefits of high resolution acquisitions are largely unexploited by traditional volume-based methods. Quantitatively, we show that the most common version of the traditional approach has spatial localization that is only 35% as good as the best surface-based method as assessed with two objective measures (peak areal probabilities and ‘captured area fraction’ for maximum probability maps). Finally, we demonstrate that substantial challenges exist when attempting to accurately represent volume-based group analysis results on the surface, which has important implications for the interpretability of studies, both past and future, that use these volume-based methods.

**Significance Statement:** Most human brain imaging studies have traditionally used low-resolution images, inaccurate methods of cross-subject alignment, and extensive blurring. Recently, a high-resolution approach with more accurate alignment and minimized blurring was used by the Human Connectome Project to generate a multi-modal map of human cortical areas in hundreds of individuals. Starting from this data, we systematically compared these two approaches, showing that the traditional approach is nearly three times worse than the HCP’s improved approach in two objective measures of spatial localization of cortical areas. Further, we demonstrate considerable challenges in comparing data across the two approaches, and, as a result, argue that there is an urgent need for the field to adopt more accurate methods of data acquisition and analysis.

## Introduction

Since the 19th century, neuroscientists have tried to relate human behavior to particular functionally specialized regions within the brain. Meaningful correlations between brain function and anatomy were first achieved by post-mortem mapping of lesion locations in subjects having specific behavioral deficits in life [Reviewed (1)]. Neuroanatomists later began subdividing the cerebral cortex into distinct areas based on cytoarchitecture [e.g., (2)] and myeloarchitecture [Reviewed (3)] with the hope that well defined brain areas could be assigned specific behavioral functions.

When noninvasive methods for mapping human brain function became available, first with PET [e.g., (4)] and then fMRI [e.g., (5)], it also became desirable to establish spatial correspondence across individuals and studies using standard stereotactic coordinate spaces, borrowing an idea from neurosurgical practice. Brodmann’s parcellation (2) became especially popular in the neuroimaging community, not necessarily because it was the best, but because Talairach and Tournoux demarcated the approximate locations of Brodmann areas in their standard space (6), which was then mapped to the population average Montreal Neurological Institute (MNI) space (7).

An early and still widely used method for assessing statistical significance in functional brain imaging, statistical parametric mapping using Gaussian random field theory, requires volumetric smoothing to satisfy its underlying assumptions [e.g. (8)], resulting in the widespread adoption of smoothing in brain imaging studies. Spatial smoothing has the seductive side effect of increasing the statistical significance of weak effects in small sample sizes, but at the expense of spatial localization precision (9, 10). Traditionally, smoothed group functional activations are then statistically thresholded and summarized by single 3D coordinates that may be assigned Brodmann’s areas or gyral and sulcal designations. Unfortunately, these standard coordinates are imprecisely related to the underlying functional neuroanatomy—the cortical areas—whose neuronal populations generate the functional activations under study (9, 10).

Besides the reductions in precision from spatial smoothing and representing brain functional neuroanatomy with single 3D coordinates, another key issue is the approach used for cross-subject alignment. Because of the high degree of individual variability in cortical folding patterns, and in the location of many areal boundaries relative to folds (11, 12), traditional volume-based methods for aligning cortical areas are imprecise across much of the cerebral cortex (13). Progress in characterizing the functions of brain areas has been impeded by these factors, along with the distributed nature of many brain functions and the lack of an accurate map of human cortical areas. Surface-based approaches enable more precise spatial localization than traditional volume-based approaches (13), and have been in use for some time, particularly in studies of the visual cortex (14-16).

The recently reported HCP-MMP1.0 multi-modal cortical parcellation (https://balsa.wustl.edu/study/RVVG) contains 180 distinct areas per hemisphere and was generated from hundreds of healthy young adult subjects from the Human Connectome Project (HCP) using data precisely aligned with the surface-based neuroimaging analysis approach (9). Each cortical area is defined by multiple features such as those representing architecture, function, connectivity, or topographic maps of visual space. This multimodal parcellation has generated widespread interest, with many investigators asking how to relate its cortical areas to data processed using the traditional neuroimaging approach. Because volume-registered analysis of cortex in MNI space is still widely used (17), this has often translated into concrete requests such as: “Please provide the HCP-MMP1.0 parcellation in standard MNI volume space.” Here, we investigate quantitatively the drawbacks of traditional volume-based analyses and document that much of the HCP-MMP1.0 parcellation cannot be faithfully represented when mapped to a traditional volume-based atlas.

There are four key differences between the traditional volume-based neuroimaging approach and the HCP-style approach (for details, see (13)): (i) *Spatial resolution of images*. The HCP acquired structural images (T1w and T2w) at 0.7 mm resolution (higher than the typical 1 mm acquisition) and used FreeSurfer’s cortical segmentation algorithms [Reviewed (18)] for robust, high-quality cortical surface models (19). For fMRI, the HCP acquired data at 2 mm isotropic resolution, better than the mean cortical thickness [2.6 mm, range 1.6 - 4 mm (13)], whereas fMRI is traditionally acquired more coarsely (typically 3 - 4 mm isotropic). (ii) *Distortion reduction*. The HCP-style approach corrects fMRI data for distortion induced by inhomogeneity in the main (B0) magnetic field using a field map, which is often neglected in the traditional approach, leading to a mismatch between the fMRI and anatomical data. (iii) *Spatial smoothing*. The HCP-style approach keeps spatial smoothing to a minimum, instead averaging within parcels when appropriate to improve statistical sensitivity and power. The traditional approach uses extensive volume-based spatial smoothing to increase statistical significance, satisfy statistical assumptions, and/or strive to compensate for imperfect cross-subject alignment. (iv) *Cross-subject alignment*. The HCP-style approach gently initializes alignment of cortical areas on the surface using cortical folding patterns and then aligns areas across subjects using ‘areal features’ (cortical myelin content, resting state networks, and resting state topographic maps) that are more closely related to areal boundaries than are structural image intensities in the volume (13) or, to a lesser extent, than folding patterns alone on the surface (20, 21). The resulting standard space is a CIFTI “grayordinates” space that contains both cortical surface vertices and subcortical gray matter voxels (19). In contrast, the traditional volume-based approach strives for alignment using linear or non-linear volume-based registration of structural image intensities to a standard volume space (Reviewed (22)]. The HCP-Style approach of surface-based alignment grew out of earlier methods that used only folding patterns (12, 23-28) and that, together with surface-based smoothing [e.g., (29)], represent intermediate approaches sharing several advantages with the HCP-Style approach over the traditional volume-based approach.

We use individual-subject cortical parcellations from the HCP-MMP version 1.0 generated by an automated areal classifier (13) as a “silver standard” (see Main Text Discussion and Supplementary Discussion Section D4) to illustrate and quantify the impact of key acquisition and analysis choices on the spatial localization of cortical areas. The individual subject parcellations were derived from each subject’s multimodal imaging data in a spatially agnostic manner^1^, and we use a validation group of subjects that share no family relationships with the subjects used for parcellation or classifier training. These steps minimize concerns about possible circularity in our analysis strategy (see Main Text Discussion and Supplementary Discussion Section D1). Projecting the individual parcellations into volume space using each subject’s own surfaces results in a voxel-wise map of each area in each subject, based on that subject’s multi-modal data. We use these maps to simulate the effects of different analysis strategies on the arrangement of cortical areas and their degree of spatial overlap. The degree to which each analysis approach differs from the HCP-style approach serves as a proxy for what would happen to any neuroanatomically organized dataset if analyzed in a similar way (e.g., an fMRI study, a structural MRI study, etc.). We also include intermediate approaches that use surface registration based on folding alone instead of areal features gently initialized by folding, or use substantial smoothing on the cortical surface. Finally, we illustrate the pitfalls of mapping results from traditional volume-analyzed group average data onto the surface. Overall, we hope that a deeper appreciation of these issues will accelerate the community’s migration away from traditional analyses and towards HCP-style analyses.

## Results

### R1. Comparing areal-feature-based surface registration to traditional volume alignment of cortical areas: probabilistic maps of cortical areas

We used binary ROIs from the classifier-based individual parcellations (9) of each of the 210 validation subjects to compute probabilistic maps of each cortical area (i.e. cross-subject averages of the 210 individual subject classifications of each area, see Supplemental Methods Section M1). For the volume-based analyses, individual cortical areas were mapped back to the volume using individual subject surfaces, reversing the process by which the data were originally brought on to the surface (see Supplemental Methods Sections M2 and M3). Figure 1 shows probabilistic maps of five exemplar areas spanning a range of peak probabilities. Each area is shown as localized by areal-feature-based surface registration (MSMAll, lower middle panel), and as localized by volume-based methods (FNIRT, parasagittal volume slices). One area (3b) has a peak probability of 0.92 in the volume (orange, red), whereas the other four have volumetric peak probabilities in the range of 0.35 – 0.7 (blue, yellow). Notably, the peak probabilities of these five areas are all higher on the surface (lower center panel, range 0.90 to 1) than in the volume, indicating that MSMAll nonlinear surface-based registration provides substantially better functional alignment across subjects than does FNIRT nonlinear volume-based registration.

**Figure 1.**
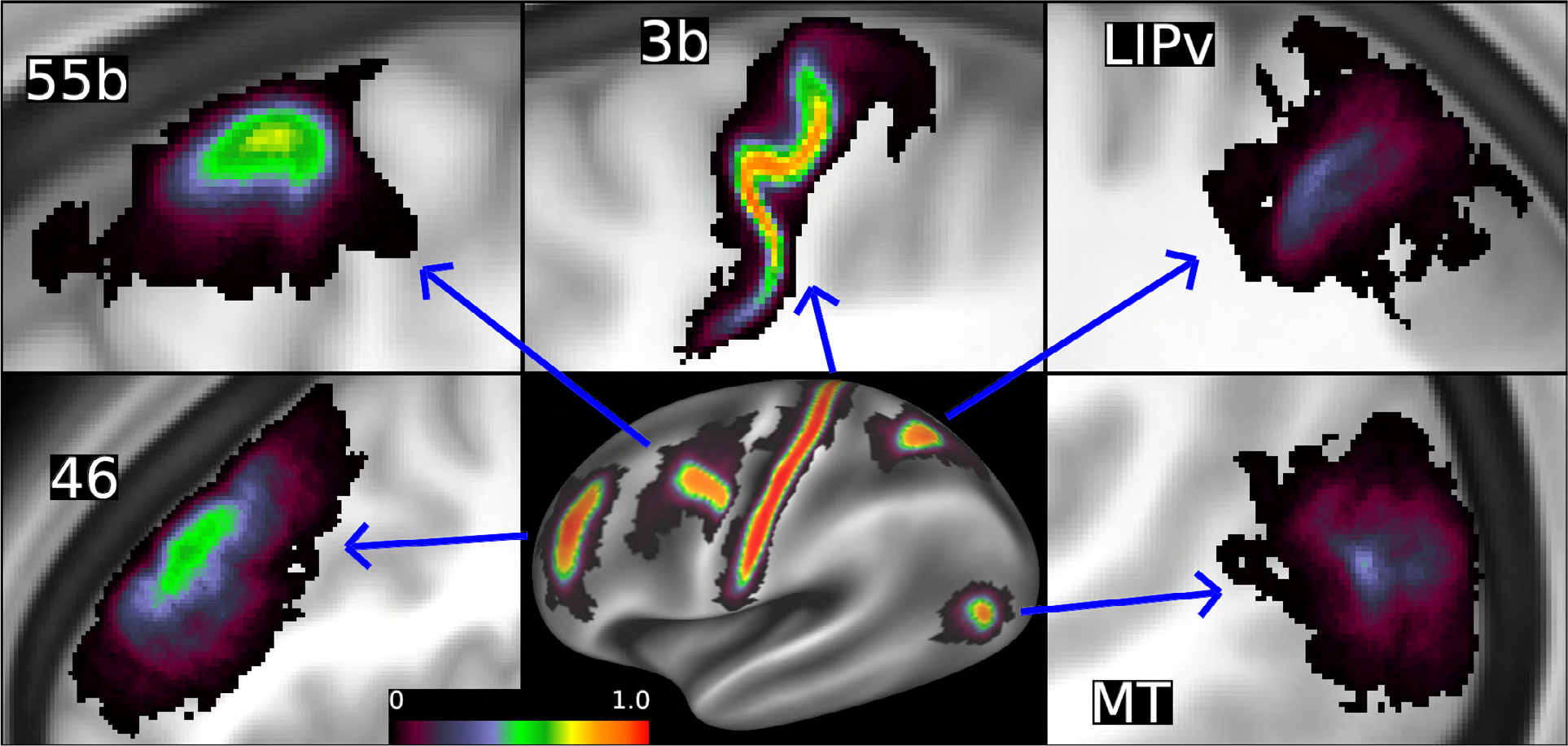
Probabilistic maps for 5 areas using both MSMAll areal-feature-based surface registration and FNIRT volume alignment. The volume-based peak probabilities are all lower than the surface-based probabilities for these example areas. Each volume-based area is shown on a parasagittal slice through the peak volumetric probability. See Supplemental Methods Sections M2 and M3. Data at https://balsa.wustl.edu/xK0Z.

In Figure 2A, the scatterplot shows that surface peak probabilities are almost exclusively higher and have many more areas with peaks at 100% (54 out of 360 on the surface versus only 3 in the volume). Peak volume probabilities have a mean of 0.70 and a standard deviation of 0.17, whereas the peak surface probabilities have a much higher mean (0.94) and a lower standard deviation (0.06). Only five of the 360 areas (R_AAIC, R_EC, L_AAIC, L_PoI1, and L_MBelt) have a higher peak value in the volume than on the surface (those below the gray line), and for these the differences are very small. Notably, most of these areas are in locations with good folding-based alignment but relatively poor fMRI SNR (which likely reduces the accuracy of MSMAll areal-feature-based alignment).

**Figure 2.**
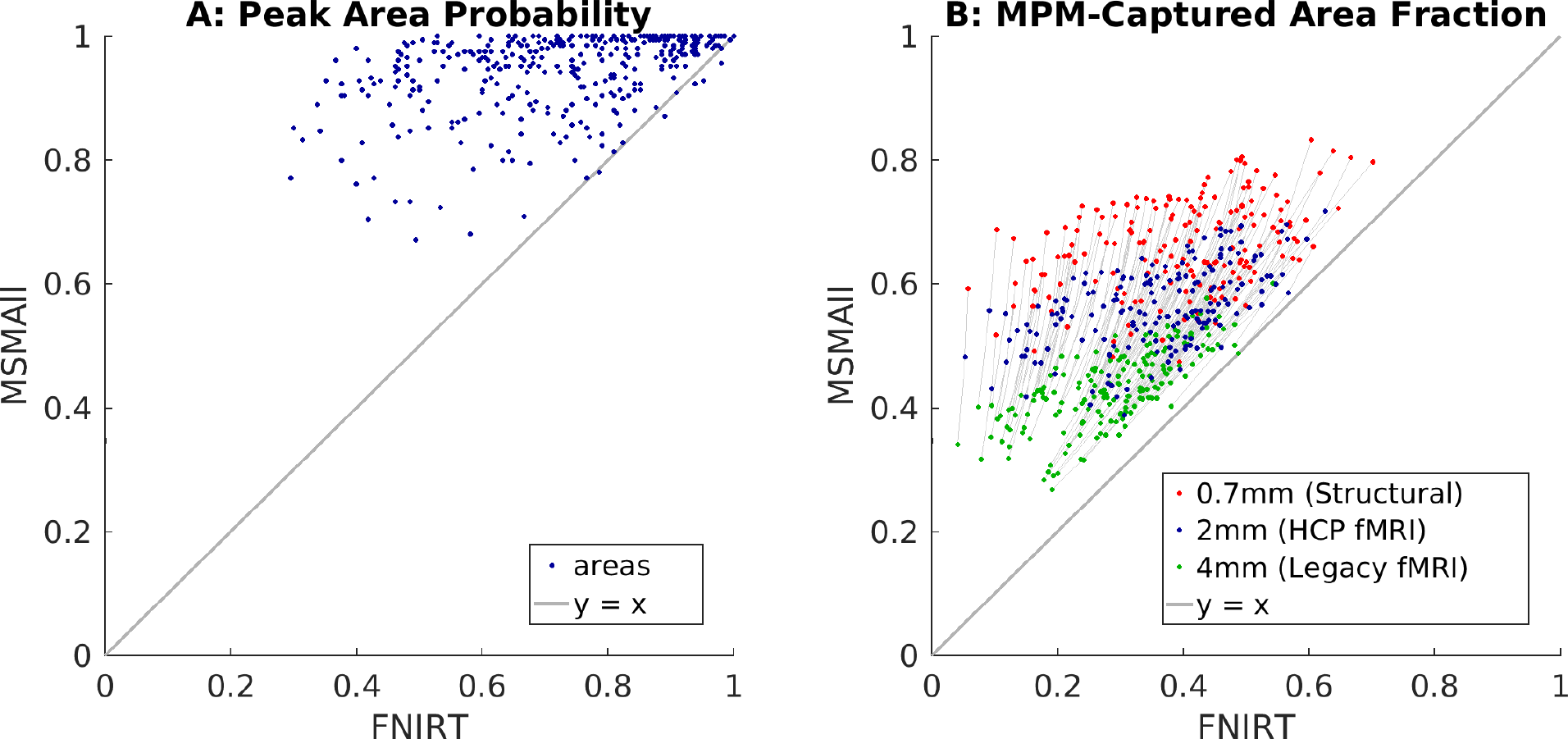
A: A scatterplot of areal-feature-based surface registration (MSMAll) peak areal probability vs volume-based registration (FNIRT) peak areal probability for all 360 areas (180 per hemisphere). B: A scatterplot of the individual areal signal captured by the group areal definitions (Maximum Probability Maps, MPMs, see Supplemental Methods Sections M6 and M7) at resolutions of 4 mm functional (e.g. legacy fMRI data, in green), 2 mm functional (e.g. HCP-style fMRI, in blue) and 0.7 mm structural (e.g. myelin or ultra high field fMRI, in red). In the right scatter plot, gray lines connect the 3 data points for each area (averaged across hemispheres, 180 total) showing the degree to which surface-based and volume-based methods benefit from increased resolution (with intermediate resolutions lying along the lines). See Supplemental Methods Sections M2, M3, M6, and M7.

The peak areal probability only represents information at a single vertex or voxel for each area. To better measure the probabilistic spatial spread of each area, we measured the proportion of each area’s vertices or voxels across all individuals that were contained within the group definition of the area, simulating the application of the group parcellation to the data (See Supplemental Methods Sections M3 and M7). This measure is derived from the idea that each cortical gray matter area will in some situations generate a distinctive ‘signal’ relative to other areas that is common across individuals (i.e. fMRI timeseries, myelin content, etc.). We use this concept for evaluating and comparing methods by asking what proportion of the individuals’ simulated signal overlaps with (or is ‘captured’ by) the surface-based and volume-based group areal definitions (from the Maximum Probability Map (MPM), see Supplemental Methods section M6). Both the surface-based and volume-based measures use the same per-individual volume files to define the simulated signal.

Figure 2B shows the fraction of the areal signal that lies within the group areal definitions, for both volume and surface. Several different resolutions are simulated, including when the data is acquired with 2 mm voxels (blue dots, simulating data acquired with HCP-style high resolution fMRI), 0.7 mm voxels (red dots, simulating data acquired at HCP-style structural resolution such as myelin maps, or ultra high field fMRI), or 4 mm voxels (green dots, simulating data acquired with “legacy” low resolution fMRI). Gray lines connect each area between its 4 mm, 2 mm, and 0.7 mm values, revealing how much each method benefits from increases in resolution. This measure is universally higher in MSMAll-aligned surface-based processing than in FNIRT-aligned volume-based processing, with very few areas even approaching equivalence. The median across areas of the surface MPM captured fraction is 0.56 for simulated 2 mm acquisition resolution, vs 0.37 for the volume MPM captured fraction. For simulated 0.7 mm acquisition resolution, the median of the surface captured fraction increases to 0.67, vs 0.41 for volume-based methods, suggesting that higher spatial resolution preferentially benefits surface-based analyses. For simulated 4 mm acquisition resolution, the median of the surface captured fraction is lower (0.43) as expected, but it remains higher than for volume-based methods (0.29), demonstrating benefits of surface-based analyses even for low-resolution, legacy fMRI data (indeed surface-based at 4mm (0.43) still outperforms volume-based at 0.7mm (0.41)). Thus, when compared with areal-feature aligned surface-based analyses, an individual’s areal signal in volume-based analyses is much more likely to be located outside of the group areal definition. Indeed on average much less than half of the signal lies inside the group areal definition in volume-based analyses, even before accounting for the smoothing that is traditionally done (see Section R4).

### R2. Comparing areal-feature-based surface registration to traditional volume alignment of cortical areas: Areal Uncertainty

We used binary ROIs of the individual parcellations from each of 210 subjects to compute maximum probability maps (MPMs) for each cortical area and for the non-cortical tissue domains (‘outside pial’ and ‘inside white’) after processing via different approaches (see Supplemental Methods Sections M1-M3, M6 and M8). As an objective measure of the quality of spatial alignment, we computed ‘uncertainty maps’, where the uncertainty value equals 1 minus the maximum probability value at each vertex or voxel. Figure 3 shows the uncertainty measure computed for areal-feature-based surface registration (MSMAll SBR Panel A) and for selected parasagittal slices of the FNIRT volume-based registration (FNIRT VBR, Panel C). The uncertainty values for MSMAll SBR are strikingly lower (better) and more consistent than those for FNIRT VBR. For MSMAll SBR, about half of cortex (49.6%) has uncertainty values less than 0.2 (maximum single area probability > 0.8), and only a small percentage of cortex has uncertainty values above 0.5 (9.0%). Note that we expect uncertainty to reach 0.5 at boundaries between two areas, and to be even higher at junctions of more than two areas, even if the registration and classification steps had resulted in nearly perfect overlap of areas across subjects. The consistently sharp transition near areal boundaries and the low overall values reflect both the high quality of areal alignment in this registration and the consistency of the parcellation across individuals. Even in regions that are typically challenging to align due to high folding variability (such as prefrontal cortex, see Supplemental Figure 1), uncertainty values near 0.5 are almost entirely confined to narrow strips along the boundaries between cortical areas. The largely uniform and low-valued uncertainty pattern, even in known challenging locations, indicates excellent cross-subject alignment with MSMAll surface registration (see Figure 6 for a comparison with surface registrations based solely on folding).

**Figure 3.**
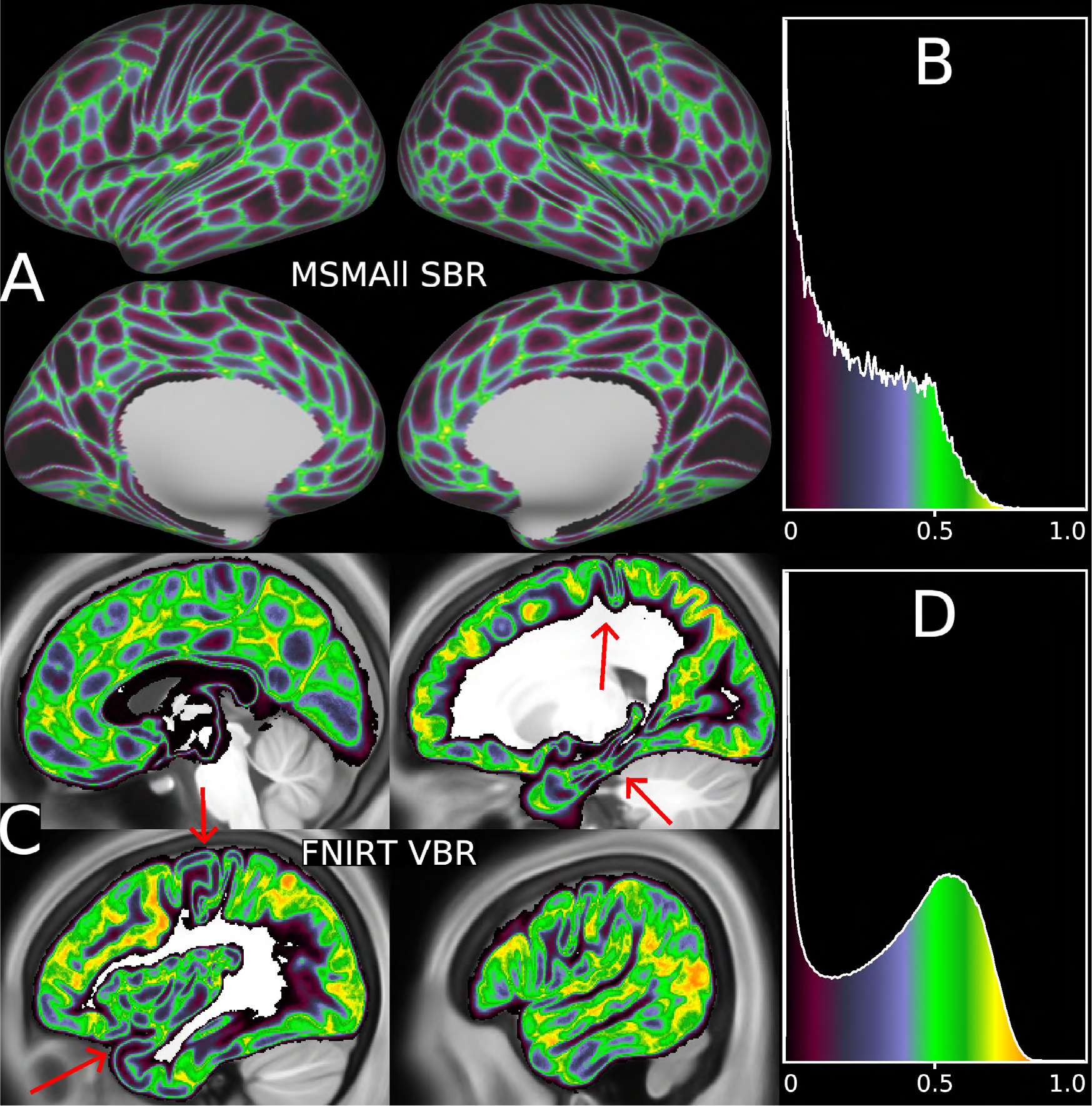
Areal uncertainty of MSMAll surface-based alignment (top) versus FNIRT volume-based alignment (bottom) for the 210V probabilistic cortical areas. The traditional volume-based approach has substantially greater uncertainty (greens, yellows, and oranges) than the HCP-Style surface-based approach as seen in the histograms (Panels B, D) as well as the images (Panels A, C). In the volume-based results, some locations have low uncertainty (purple and black) and relatively sharp boundaries between areas (red arrows: early sensorimotor, insular, and inferior temporal cortex), comparable to what is consistently found on the surface. The volume ROIs that were used to create this figure were generated by mapping the individuals’ parcellations to the 0.7 mm MNI template space using the native resolution MNI space individual surfaces and the ribbon mapping method (19). Using 0.7 mm voxels minimizes the effects of voxel size on the group probability maps, allowing the effect of alignment to be investigated separately from the effect of voxel resolution. In practice, typical fMRI resolutions lead to increased signal mixing between areas and non-cortical tissues, for both surface and volume analysis (see Results Section R6). See Supplemental Methods Sections M2, M3, and M8. Data at https://balsa.wustl.edu/PGX1.

In the FNIRT volume-based alignment, some low folding variability locations show reasonably good cross-subject agreement, such as the central sulcus and insula (red arrows). In these locations the higher uncertainties are locally restricted to obvious boundaries between areas, or between cortex and the CSF or white matter. However, very little cortex has uncertainty below 0.2 (only 3.3% of the voxels where cortex is the most likely tissue), and almost two thirds of cortex has uncertainty above 0.5 (65.5% of voxels where cortex is the most likely tissue). The higher uncertainty is concentrated in regions of high cortical folding variability (See also Supplemental Figure 1). Additionally, the volume-based uncertainty maps contain both uncertainty in gray matter alignment and uncertainty in areal alignment (See also Supplemental Figure 2). These high uncertainties show that volume-based registration failed to align the HCP-MMPv1.0 areas in many locations. Notably, most of the very low values in the volume histogram (Figure 3D) are from the wide low-uncertainty fringes that are in atlas white matter and CSF, rather than in locations that are highest probability gray matter (see Supplemental Figure 3). In contrast, the low values of the surface uncertainty (upper right panel) occur exclusively inside gray matter cortical areas. Altogether, this indicates that FNIRT-based volume analysis is unable to reliably discriminate between cortical areas over much of the neocortex.

### R3. Volumetric Areal Maximum Probability Maps (MPMs)

Volumetric MPMs for cortical areas have been reported in other studies (e.g., (30)), and we generated volumetric MPMs for the HCP-MMP1.0 parcellation, as described in Supplemental Methods Section M6. We found that in regions where the probabilistic gray matter ribbon has relatively high values and low areal uncertainties, the vMPM forms a thick continuous ribbon, roughly comparable to average cortical thickness in these regions. In such regions, volume-based alignment is not at a major disadvantage to surface-based alignment. In contrast, for regions where probabilistic cortical gray matter is less well aligned and areal uncertainty is consistently high the vMPM is thinner than the average cortical thickness. Indeed, in a few locations there are overt gaps that lack a winning cortical area, identifying regions where white matter or CSF is more likely than any single cortical area (see left side of Supplemental Figure 4). This contrasts with the accurate alignment of each individual subject’s parcellation, mapped to the volume using the subject’s surfaces and displayed on the individual’s T1w volume, which completely overlaps the map of the individual’s gray matter (see right side of Supplemental Figure 4). More generally, the volumetric probabilistic maps for the exemplar areas shown in Figure 1 represent the expected distribution of data generated by these areas in any dataset that has been registered using FNIRT using the HCP’s FNIRT configuration without spatial smoothing. The net result is that each area in the vMPM is much smaller than its corresponding probabilistic map, such that a large fraction of each area’s group probability (and therefore signal) will fall outside the vMPM parcel. We quantified this effect above in Figure 2 for MSMAll surface-based registration vs FNIRT volume-based alignment and below in Figure 8 for additional analysis approaches. This poor alignment of individual subject cortical areas to the group MPM is a fundamental problem for using a volumetric MPM to represent cortical areas. We next demonstrate that this problem is dramatically exacerbated by the spatial smoothing that is commonly used in volume-based studies.

**Figure 4.**
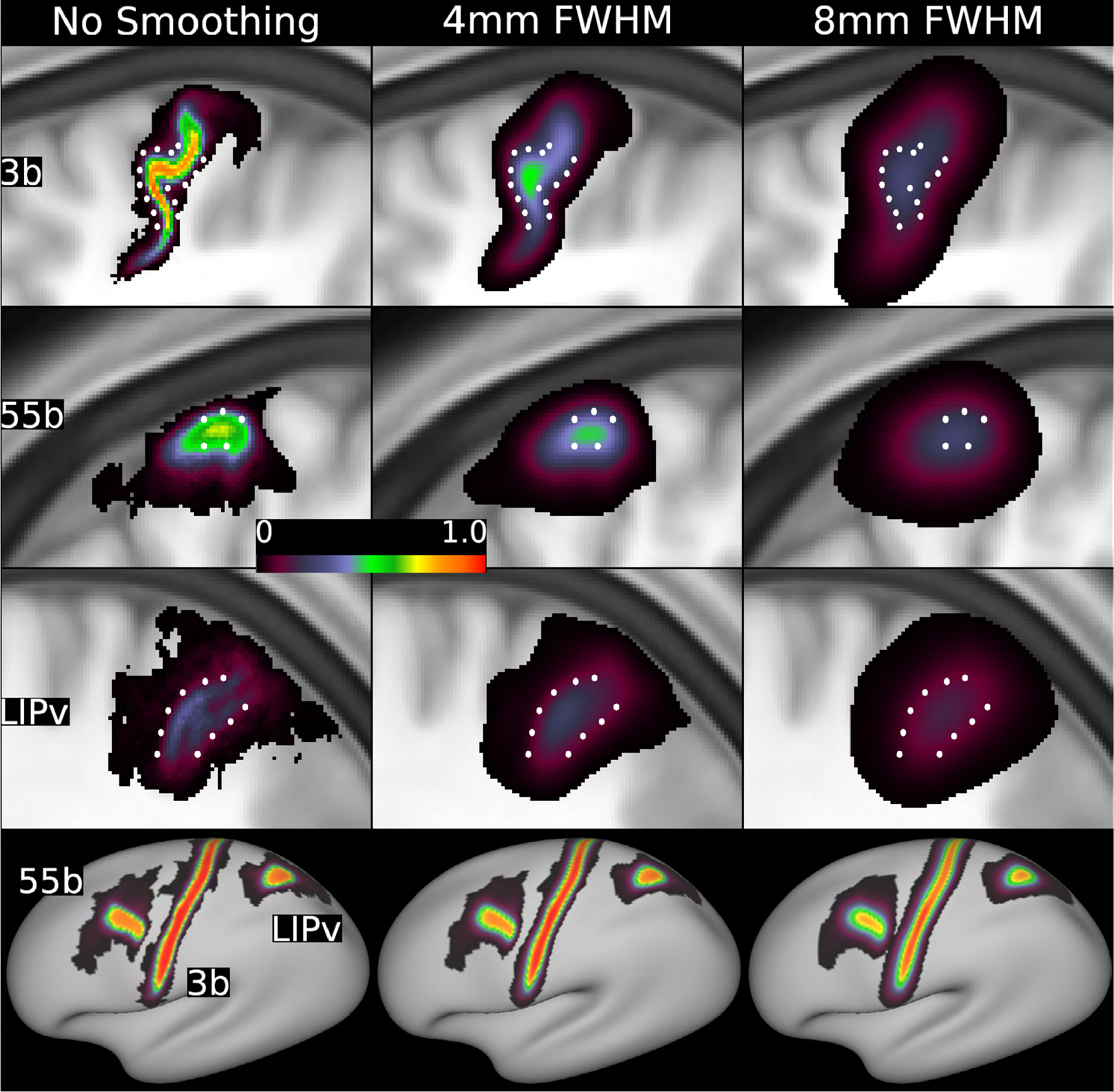
Effects of volume-based and surface-based smoothing on example cortical areas. The top three rows show enlarged sagittal slices of volumetric probabilistic maps through the maximum probability of 3 exemplar areas, before (left) and after unconstrained volume-based Gaussian smoothing of 4 mm (center) or 8 mm (right) FWHM. In each row, white dots are in corresponding positions for reference. The bottom row shows the same amounts of surface-based Gaussian smoothing applied to the same 3 areas after areal-feature-based registration (MSMAll). Areal probability values decrease substantially more in the volume after smoothing than on the surface with the same amount in millimeters of smoothing. See Supplemental Methods Sections M3 and M4. Data at https://balsa.wustl.edu/7Blg.

### R4. Effects of Spatial Smoothing in the Volume and on the Surface

Smoothing of volumetric data is widely used as a way to reduce voxel-wise noise, increase measures of statistical significance, and satisfy statistical assumptions. It is also often presumed to compensate for imperfect alignment of cortical areas across subjects. Unfortunately, smoothing in the volume mixes data across tissue compartments and across areal boundaries, including non-contiguous areas on opposite banks of gyral and sulcal folds (13). By treating the binary individual-subject parcellations as patches of idealized signal, we can show the effect of smoothing on the purity and extent of areal signal (See Supplemental Methods Section M3 and M4). This effect is visible as a reduction in peak areal probability and an expansion of the volume-based areal probability maps in Figure 4, which shows three exemplar areas after different smoothing amounts. Area 3b (top row) has a relatively tight probabilistic distribution without any smoothing (left), but the peak value is markedly reduced and the spatial extent increased with 4 mm FWHM volumetric smoothing, deleterious trends that worsen with 8 mm FWHM smoothing. Areas 55b and LIPv start out with greater spread in the no smoothing condition, so the effects of volumetric smoothing are not as visually dramatic, but they are nevertheless substantial in both cases. Comparable levels of surface-based smoothing applied to the same three MSMAll-registered areas (bottom row) show a much smaller effect, though smoothing still erodes localization.

From the standpoint of cortical localization, volume-based smoothing substantially erodes the fidelity with which areal assignments can be made. This effect is illustrated in the top two rows of Figure 5, which shows areal uncertainty maps (second row) and histograms (top row) without smoothing (left) and after volumetric smoothing of the group probability maps by 4 mm FWHM (middle column) and 8 mm FWHM (right column), which are commonly used levels of volumetric smoothing in fMRI studies. Over most of the cortical ribbon, areal uncertainty in the volume-smoothed maps exceeds 0.5 (green/yellow), especially for 8 mm FWHM smoothing, indicating that neuroanatomical identification at the level of individual cortical areas in the HCP-MMP1.0 parcellation is quite limited indeed. Surface-based smoothing of the areal probability maps at 4 mm and 8 mm FWHM (Figure 5, bottom rows) also causes some blurring of areal boundaries. However, unlike volume-based smoothing, it does not blur across sulci or across tissue categories, so the overall effects are substantially less deleterious. See Supplemental Figure 5 for additional volume smoothing levels of 2 mm, 6 mm, and 10mm FWHM that have been reported for fMRI studies (17), and Supplemental Figure 6 for the additional surface smoothing level of 15 mm FWHM that has been reported in the literature (31) and that approaches the areal uncertainty values seen in unsmoothed volume-based alignment and the same three levels of smoothing (4 mm, 8 mm, and 15 mm) with a FreeSurfer alignment.

**Figure 5.**
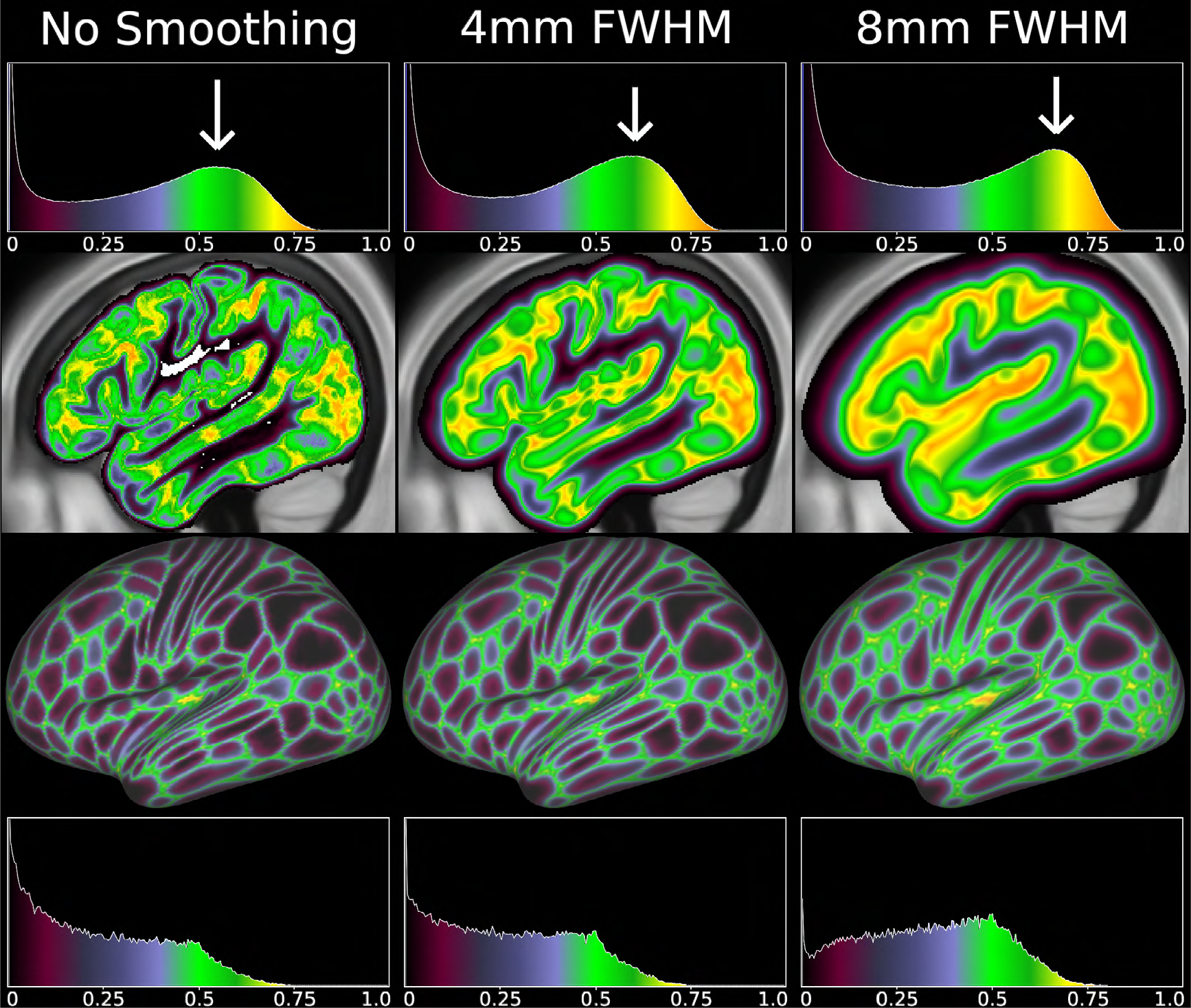
Comparison of different degrees of smoothing (columns) for both volume-based (top two rows) and surface-based (bottom two rows) approaches. Both areal uncertainty maps and histograms are shown. These were computed by smoothing the probability maps, which is equivalent to smoothing the per-subject ROIs before averaging. Smoothing kernels on the surface clearly have less deleterious effects than smoothing kernels of the same size in the volume, because surface smoothing avoids smoothing across sulci or into other tissues. As with Figure 3, the volume-based histograms have substantial “low uncertainty” tails that arise from poor alignment of the cortical ribbon, and from the tail of the Gaussian smoothing kernel within the white matter and CSF. See Supplemental Methods Sections M3, M4, and M8. Data at https://balsa.wustl.edu/6MB7.

Volume-based smoothing also shifts the location of maximum gray matter probability towards the central CSF-filled portion of sulci and towards the white matter portion of gyri (Supplemental Figure 7). Features in regions of low folding variability (e.g., insular cortex and the superior temporal gyrus) are sharply defined without smoothing, as is the boundary between gray and white matter in such regions. With smoothing, the sulcal fundi are not only blurry, but the apparent location of the transition between gray and white matter is substantially shifted, particularly for high smoothing levels (e.g. 8 mm FWHM).

### R5. Comparing areal alignment quality of different surface-based registration methods

We compared the alignment quality of four different surface-based registration methods (See Supplemental Methods Section M4). Figure 6 shows results for MSMAll registration (areal feature based), MSMSulc (folding based, with less distortion than FreeSurfer), FreeSurfer (folding based), and a sphere rotation-only rigid alignment (which has only 3 degrees of freedom, whereas spherical warpfields often have thousands) derived from the FreeSurfer registration. Each row shows surface probabilistic maps for six representative cortical areas, areal uncertainty maps, and histograms of uncertainty values. All three ways of representing the data demonstrate that MSMAll provides the tightest alignment of HCP-MMP1.0 probabilistic maps (higher peak probabilities and tighter clustering). MSMSulc is intermediate, though only slightly better than FreeSurfer, which in turn is slightly better than spherical rotation overall. The spherical rotation method’s alignment is driven primarily by similarities in spherical inflation across subjects. The peak areal probabilities and degree of areal dispersion found here for FreeSurfer folding-based registration is comparable to that reported in previous studies that used FreeSurfer registration with other parcellations (12, 32, 33). Note that the maximum possible overlap on the surface (or in the volume) is less than 100% for some areas because the areal classifier does not detect 100% of all areas in all subjects and because there are atypical topologies in some areas in some subjects that prevent any topology preserving registration from achieving perfect overlap across subjects (see Supplemental Methods Section M1).

**Figure 6.**
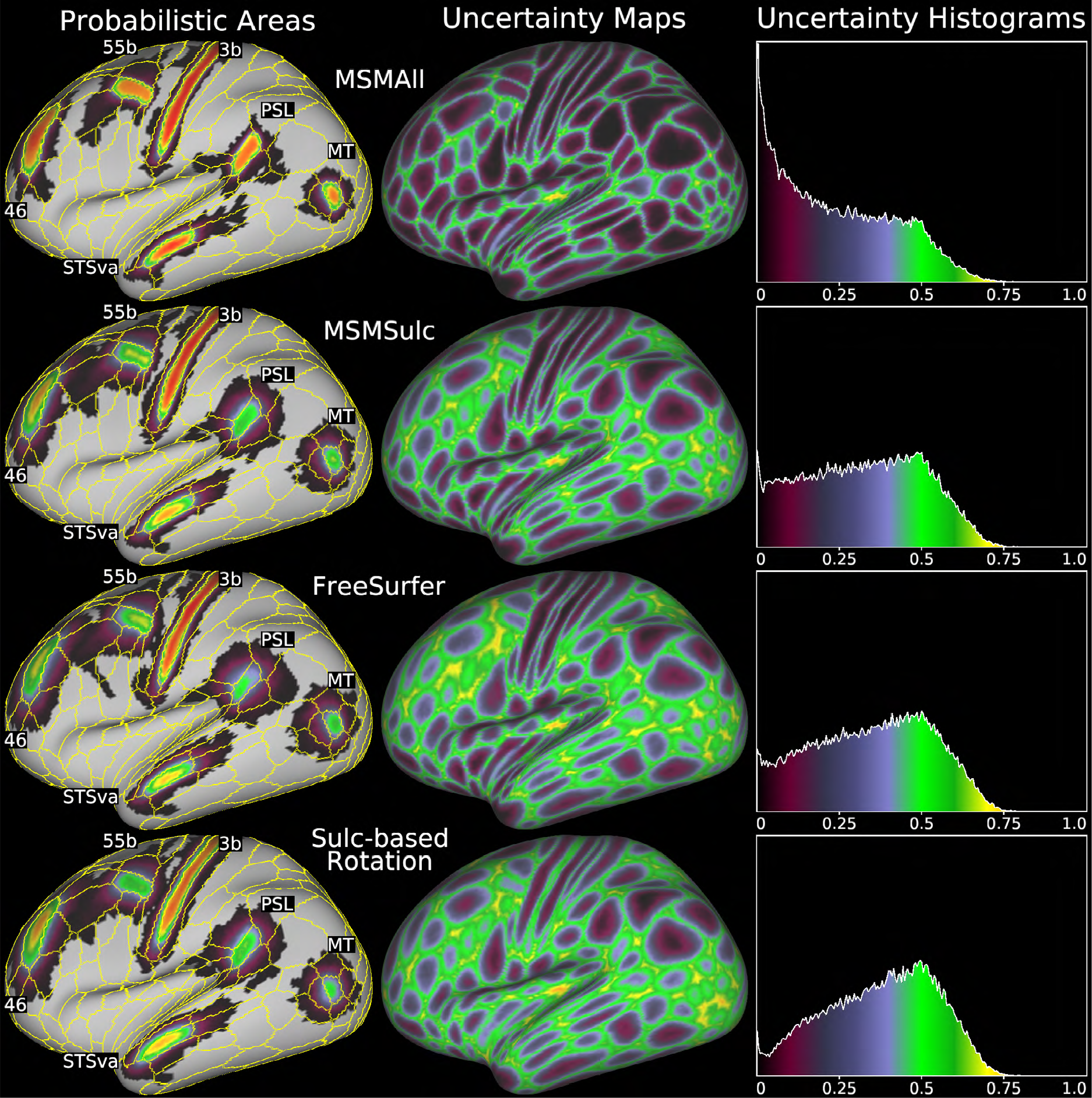
Comparison of four surface-based alignments: MSMAll areal-feature-based registration (top), MSMSulc folding-based registration (second row), FreeSurfer folding-based registration (third row), and a rigid spherical rotation alignment based on the FreeSurfer registration (bottom). The left column shows six probabilistic areas for each registration approach with yellow contours representing the areal boundaries from the 210V MPM. The center column shows the maps of areal uncertainty (1 – maximum probability at each vertex), as in Figure 3. The right column shows the histograms of the uncertainty maps. See Supplemental Methods Section M4. Data at https://balsa.wustl.edu/1616.

### R6. Effect of Acquisition Resolution

Of the three main acquisition and analysis choices made in neuroimaging studies that directly impact spatial fidelity (acquisition resolution, method of cross-subject alignment, and method and amount of smoothing), we found that commonly used fMRI acquisition resolution choices have the smallest overall impact (See Supplemental Methods Section M2). Figure 7 compares how surface-based and volume-based processing is affected by the combination of partial-volume and volume-based tissue alignment effects, at current state-of-the-art 3T fMRI acquisition resolutions (2 mm isotropic) and five other simulated resolutions (0.7 – 4.0 mm). It shows that partial-volume effects on the surface decrease substantially (gray matter signal fraction increases) when acquiring data with smaller voxel sizes. Notably, for surface-based analysis, the group-average spatial pattern in this measure is largely determined by cortical thickness, and is highly uniform over much of the cortex. In contrast, the histograms of group average volume data are largely unchanged despite increases in acquisition resolution, because the inaccuracies of volume-based alignment largely dominate the measure, showing that traditional volume-based analysis is unable to fully take advantage of higher resolution data. The maximum volume-based group cortical signal fraction also varies considerably across different regions of cortex (e.g. between the central sulcus and superior parietal cortex), suggesting spatial heterogeneity in statistical power and localization precision for volume-based analyses.

**Figure 7.**
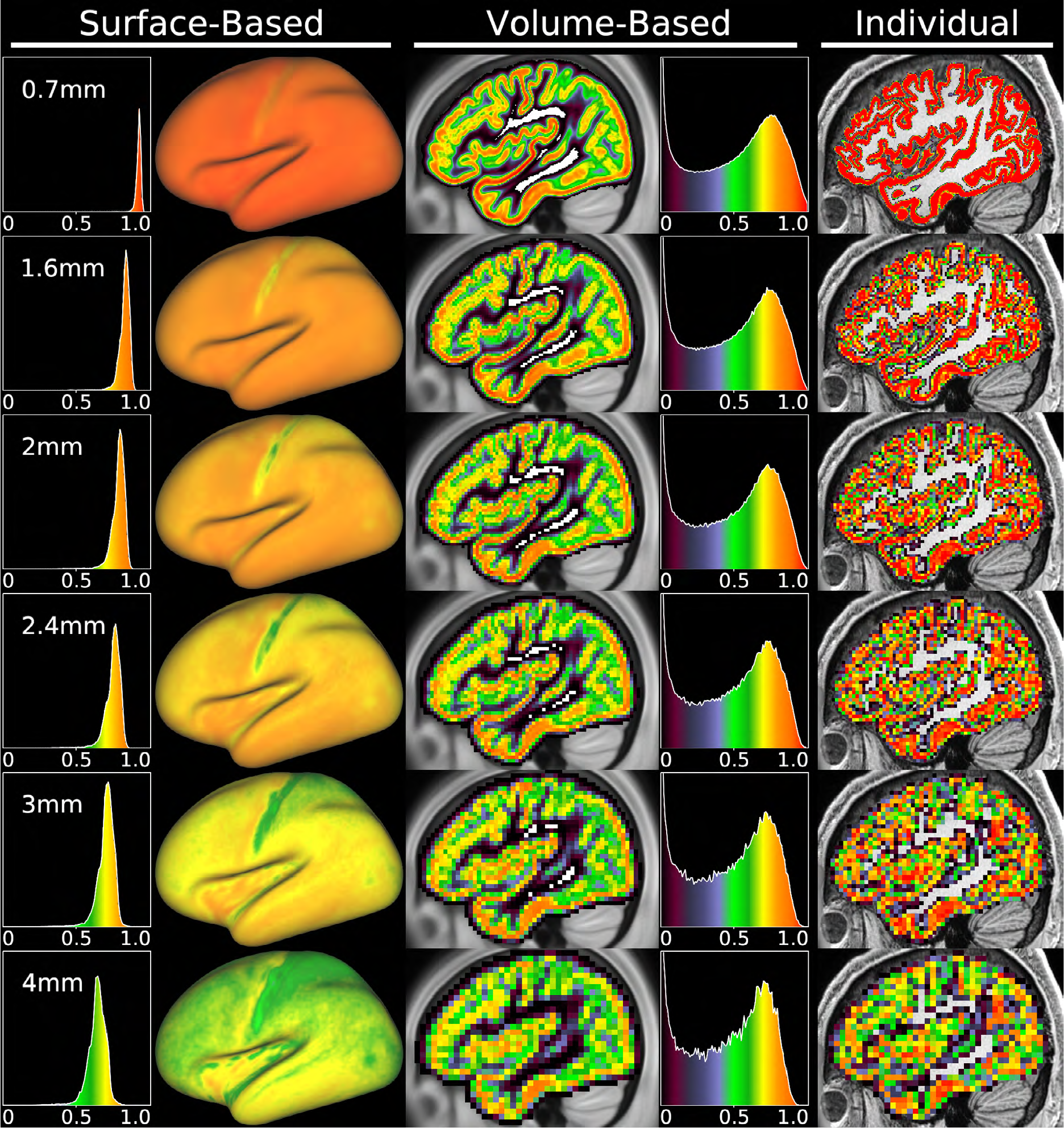
shows the effect of acquisition resolution on the separation of cortical signal from non-cortical signal, for surface-based (left two columns) and volume-based (next two columns) processing. The measure shown is the group average cortical gray matter fraction of each vertex or voxel. The rightmost column shows an individual’s (HCP subject 121618) cortical fraction volumes for the same six resolutions, as an example of the inputs to the analyses. Smoothing was not applied. The cortical signal fraction becomes somewhat degraded at the edge of cortex (green voxels) in many regions even at 2 mm resolution (even though this is less than the mean cortical thickness) and is severely degraded (many green and blue voxels) at traditionally used resolutions between 3 and 4 mm. See Supplemental Methods Section M2. Data at https://balsa.wustl.edu/5gMx.

Although acquisition resolution has the lowest impact among the three aforementioned processing choices, finer acquisition resolution, especially below the mean cortical thickness of 2.6 mm, is very helpful for surface-based studies, which are not limited by volume-based cross-subject alignment. Higher resolutions (e.g., the 1.6 mm voxel size for HCP 7T fMRI data) will reduce partial volume effects and enable more advanced analyses, such as those focusing on laminar differences within the cortical ribbon (9) (see Supplemental Figure 8). However, such analyses will require technical advances in MRI acquisition, and optimization of tradeoffs between voxel size, signal-to-noise ratio (SNR), and acquisition speed.

### R7. Summary localization measures for different registrations and smoothing levels

Figure 8 provides a valuable summary comparison across a variety of approaches, using the aforementioned peak probability and ‘area capture’ measures (see Figure 2) for each of the 360 cortical areas. Notably, this figure also includes a new strain-regularized MSMAll surface-based registration (20), which was not used in defining the parcellation or training the areal classifier, but nevertheless shows very similar performance to the original MSMAll. The left panel box and whisker plots show the peak probabilities of each area, and the right panel shows the MPM captured area signal measures for the same ten methods (using 2 mm resolution partial volume weighting). These measures give the same ranking to the medians of each method, which are tabulated in Supplemental Table 1. A particularly important comparison shows that the most commonly used smoothing level (17) in the traditional approach (FNIRT + 8 mm FWHM volume smoothing) is only 35% as good as the best surface-based method (MSMAll) using measures of median peak areal probability (0.340 vs 0.967) and median MPM-captured areal fraction (0.200 vs 0.564). Comparing the surface-based analysis using only rigid rotation for spherical alignment to the volume-based analysis without smoothing reveals the benefits achieved by simplifying the more challenging cross-subject cortical registration problem from 3D to 2D and solving the more tractable tissue segmentation problem to handle the third dimension. These conceptual improvements reflect the fundamental advantage that surface-based approaches have over volume-based approaches (see Supplementary Discussion Section D2). Large amounts of surface-based smoothing (15 mm FWHM) substantially degrade cortical area localization to levels similar to volume-based alignment with no smoothing.

**Figure 8.**
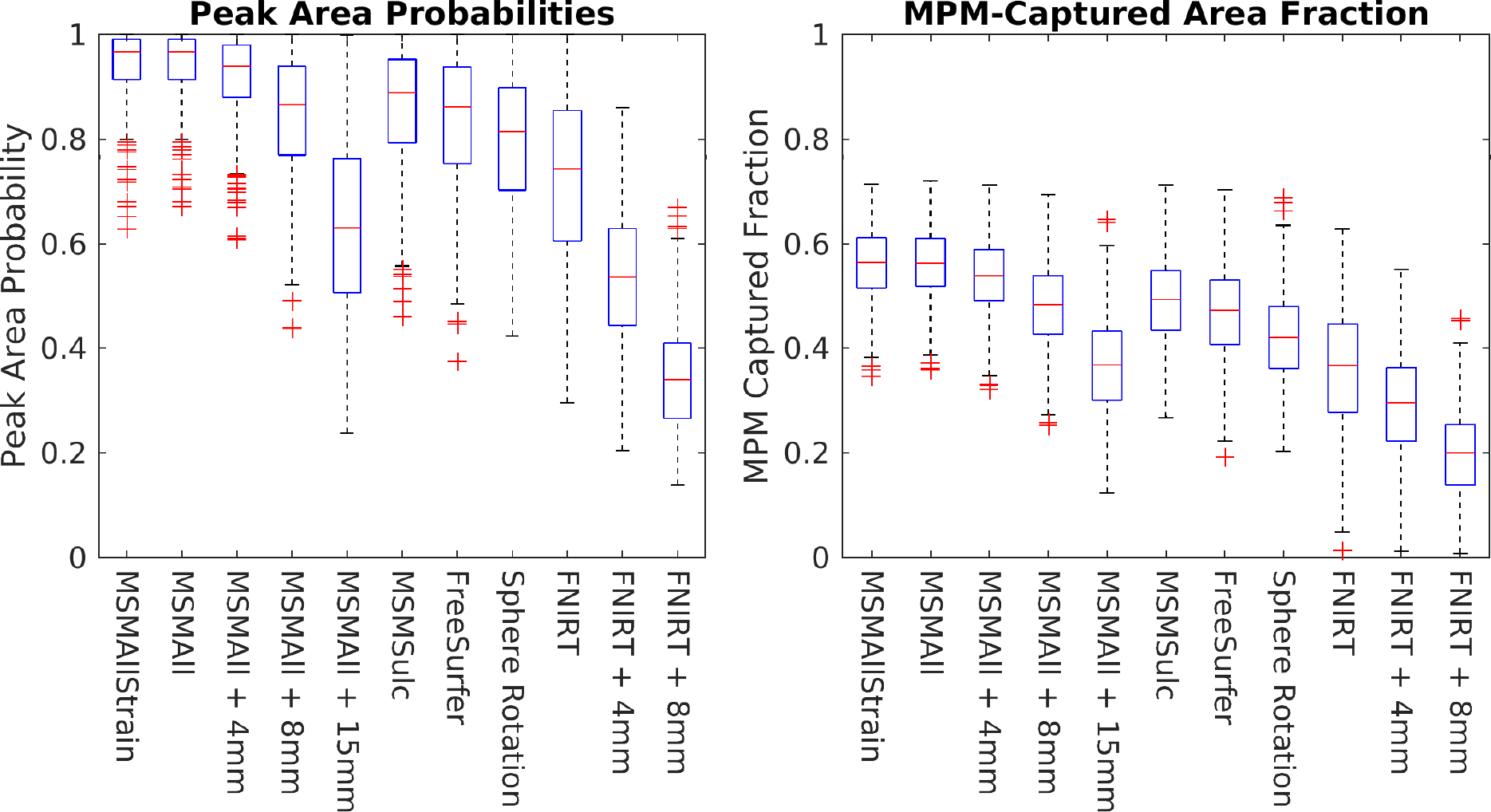
Left: Box and whisker plots of the peak probability of each area for various SBR methods and for FNIRT volume-based registration, plus the effects of differing amounts of surface (4 mm, 8 mm, and 15 mm FWHM) and volume smoothing (4 mm and 8 mm FWHM). Less optimal registration methods and greater smoothing consistently reduce peak areal probability. Volume-based smoothing has the largest impact, followed by volume-based versus surface-based alignment. The decrease of FreeSurfer compared to MSMSulc is similar in magnitude to that of smoothing MSMAll data by 4mm FWHM. Right: MPM captured area fraction using 2 mm MNI space voxels for the same ten methods, showing a similar pattern. Notably, the areas that do worse in the new MSMAllStrain are generally well aligned by folding, whereas the areas that do better in MSMAllStrain have more variability across subjects (the new MSMStrain allows more mild to moderate distortions while clamping peak distortions). Red line is the median, box edges are the 25 to 75 percentiles, whiskers are 2.7 standard deviations, and red pluses are outliers beyond 2.7 standard deviations. See Supplemental Methods Sections M2-M7.

### R8. Mapping legacy group average volume results onto the surface

Traditional volume-based analyses often map group average volume-based results onto group average surfaces for visualization purposes, using, for example, the ‘average fiducial mapping’ approach (24). While this approach has known limitations, its accuracy has not previously been analyzed carefully. We used a modified form of this approach, which we call ‘average surface mapping’ (ASM), employing the ribbon-based volume to surface mapping technique (19) and the group average MSMAll white and pial surfaces (See Methods Section M9). In Supplemental Figure 9 we illustrate the primary pitfall of this approach: Group average surface contours do not consistently follow the group average cortical ribbon, particularly in regions of high folding variability (see panel B2). Even when using folding-based surface registration, topologically incompatible folding patterns (e.g., two gyri in some subjects where there typically is only one) lead to reductions in group average cortical surface area that “shrink” the surface towards the direction of folding, as these patterns are unable to be aligned and therefore result in significant cross-subject variability of coordinates. The folding detail of the group average surface is further reduced when using average surfaces from MSMAll-registered data instead of folding-based alignment because of discrepancies between function and folds (see Supplemental Figure 1). Indeed, the MSMAll group average surfaces only show significant folding detail in locations that have good correspondence between folds and areas. Thus, mapping group average volume data onto group average surfaces will have additional biases (on top of the blurring effects from misalignment and smoothing and the biasing effects of smoothing folded cortex shown in Supplemental Figure 7). Folding-based average surfaces will be only modestly better that MSMAll surfaces overall.

An alternative approach for mapping group average volume data to surfaces is the ‘multi-fiducial mapping’ method, using anatomical midthickness (fiducial) surfaces from many individuals as intermediates (24). Here we similarly modified this method by using ribbon mapping and call this ‘multiple individual mapping’ (MIM). The cortical gray matter fraction map from this approach is smoother, showing less sensitivity to folding patterns, but also has a lower overall value, as shown in Supplemental Figure 9 (see Figure Legend and Supplemental Methods Section M2 and M9). This effect occurs because the tissue misalignment from FNIRT is applied twice, once in making the volume-based group average cortical ribbon, and again when mapping the group average onto the individuals’ surfaces. This method also results in more mixing between tissue classes, decreasing the cortical contribution to the surface-mapped values.

These effects are apparent when looking at cortical areas as well. Indeed, after averaging the 2 mm MNI-space area maps in the volume, mapping this result onto a large set of individual surfaces, and averaging on the surface, the resulting area maps are dramatically changed relative to the surface-based approach of mapping each individual subject’s area volumes onto their own surfaces before averaging (Supplemental Figure 10). These effects extend to the maximum partial volume maps as well (Figure 9). In regions with high folding variability, it is challenging for cortical areas to be dominant over non-cortical tissue classes (white matter in particular), as shown by the extensive bright yellow regions in the top two rows for FNIRT + ASM mapping (column 2) and their even greater extent for FNIRT + MIM (column 4). These effects are further exacerbated by volume-based smoothing (columns 3 and 5). Notably, for some cortical areas that are well aligned by folds, such as those in the insula, the methods are essentially identical across unsmoothed approaches (though again, volume-based smoothing is universally deleterious). However, when analyzing all of cortex, it is much better to map individual data onto individual surfaces and align the data on the surface if one wants to relate it to surface-based data, including the HCP’s multi-modal parcellation.

**Figure 9.**
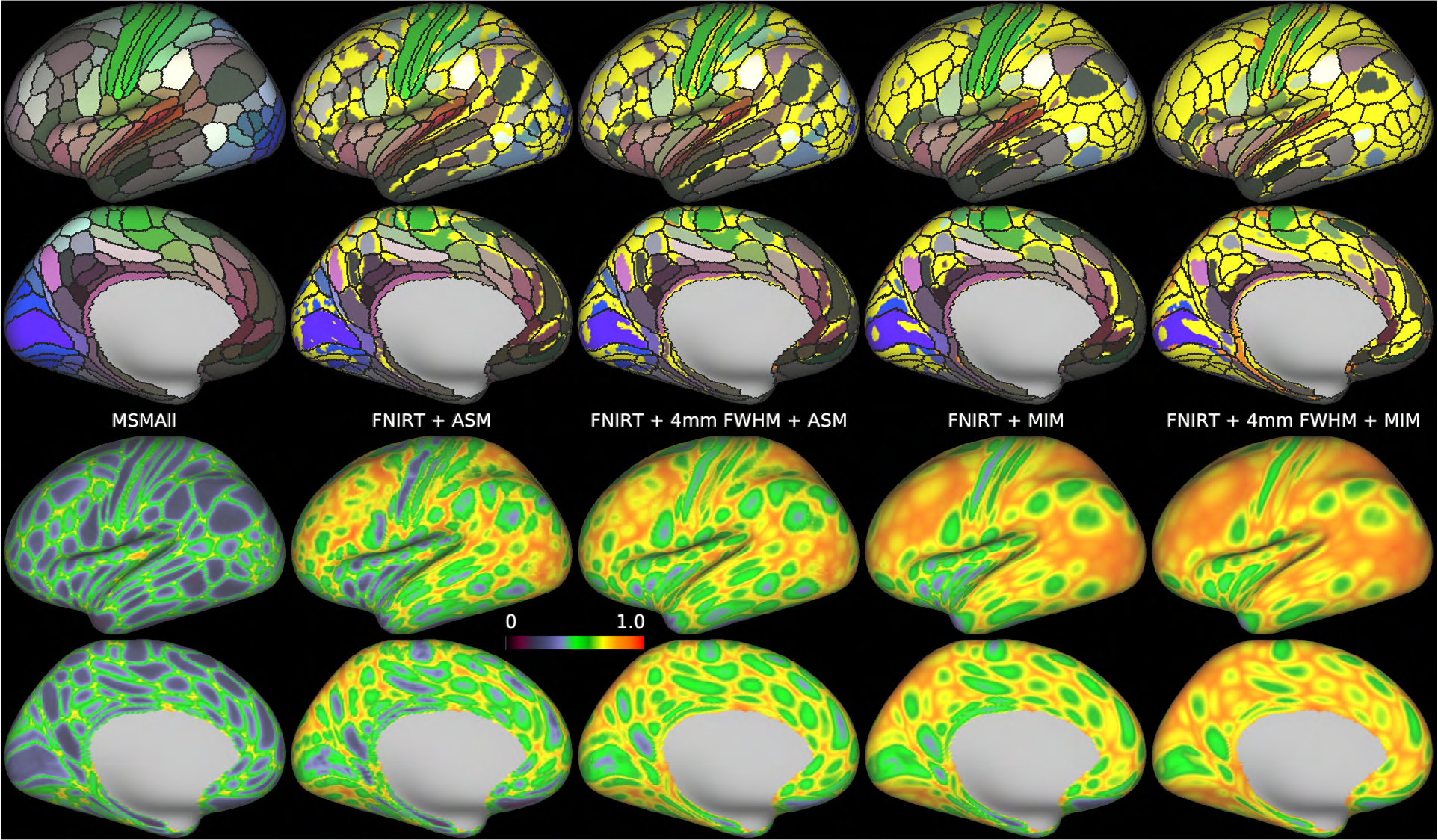
Comparison of the surface-based maximum partial volume map to the maps produced after volume-based analysis with Average Surface Mapping (ASM) or Multi-Individual Mapping (MIM), and 4 mm FWHM volume-based smoothing before ASM and MIM. The figure uses the same methods as Supplemental Figure 10, and then uses the maximum fraction to label the surface vertices. In the top two rows, bright yellow is the white matter label, and bright orange is the CSF label (occurring in only a few small patches). Substantial regions of the cortex are not separated into cortical areas after volume-based analysis and MIM, and ASM shows significant stripes where the gyral crowns are decapitated. On the other hand, in regions of lower folding variability and variability of areas vs folds such as the insula, volume-based methods reproduce the parcellation found with the surface-based approach, particularly if smoothing is not used. See Supplemental Methods Sections M2 and M9. Data at https://balsa.wustl.edu/nKvx.

## Discussion

We have systematically explored the impact on spatial localization of different acquisition and analysis choices, using a large dataset in which the cerebral neocortex was individually parcellated using multiple modalities. This data allowed us to incisively and quantitatively assess the impact of these methodological choices on spatial localization, not only for the HCP-MMP1.0 parcellation data but also by extension for a wide range of neuroimaging studies (both past and future). The analysis in Figure 8 summarizes the key results and illustrates the magnitudes of the effects on spatial localization of different analysis choices. Areal-feature-based surface registration (MSMAll or MSMAllStrain) without smoothing clearly achieves the best localization of the HCP-MMP1.0 areas. The effect of 4 mm FWHM surface-based smoothing on MSMAll registered data is comparable to the difference between MSMSulc and FreeSurfer folding-based registrations: modest, but still meaningful. The difference between registrations based on areal features vs folding is comparable to the difference between folding-based surface registration and FNIRT volume-based registration, and also similar to the difference between unsmoothed areal-feature-based surface registration and more heavily smoothed areal-feature-based surface registration at 8 mm FWHM. 15 mm FWHM surface-based smoothing more substantially degrades MSMAll aligned cortical areal spatial localization to be similar to that of unsmoothed FNIRT volume-based alignment. The degradation of spatial localization from moderate (4 mm FWHM) unconstrained volume-based smoothing is similar to the difference between unsmoothed volume-based registration vs areal feature-based registration, and the degradation is substantially greater for the more commonly used 8 mm FWHM volume-based smoothing (17). Importantly, these effects are cumulative, and one cannot “recover” any spatial localization that was lost in previous steps (i.e. smoothing does not ameliorate reductions in spatial localization from registration methods that do not align areas, it instead makes them worse).

Studies that use extensive smoothing typically do not overtly justify their chosen amount of smoothing. The most common smoothing level in the literature is 8 mm FWHM, the default value in SPM, the most commonly used neuroimaging software package (17). While spatial smoothing is indeed effective in increasing statistical sensitivity, so too is improving areal alignment across subjects with surface-based, areal-feature-based registration. Aligning “like with like” tends to improve z-statistics by reducing cross-subject variability (see (13)), and it also makes the resulting group maps sharper and more neuroanatomically interpretable rather than making them blurrier and less interpretable as smoothing does. Use of areal feature-based registration together with minimal smoothing was critical for creating HCP’s multi-modal parcellation at the group level (9). Permutation-based non-parametric statistical methods (such as that offered in PALM software [https://fsl.fmrib.ox.ac.uk/fsl/fslwiki/PALM; (34)] can be used without smoothing, or with small amounts of smoothing, in contrast to popular parametric methods, and also work well with HCP-Style CIFTI or parcellated data. The statistical assumptions required by Gaussian random field theory, which have been assumed to be satisfied by using large smoothing kernels, were recently shown to be unsatisfied in some cases, leading to increased false positives (35).

Surface-based representations of cortical data are more visually informative than volume-based depictions, insofar as the convoluted sheet-like cerebral cortex is best inspected on a 2D visual model, often using inflation or flattening to expose cortex that is buried inside sulci. Modest surface-based smoothing (e.g., 4 mm FWHM) can increase statistical sensitivity and make maps more visually appealing at the cost of a modest decrease in spatial resolution (e.g., reducing the discriminability of thin features such as area 3a or fine details such as somatotopic variations in myelin content in areas 4 or 3b). Parcellation provides a more powerful alternative to spatial smoothing for increasing statistical sensitivity (13). Parcel-constrained averaging can be considered a neuroanatomically informed method of “smoothing” (averaging within cortical areas defined using multiple modalities)^2^ that also greatly reduces the number of statistical comparisons. Rather than improving alignment, spatial smoothing further erodes the ability to localize brain structure, function, or connectivity in neuroimaging studies. The statistical sensitivity benefits of spatial smoothing come from reducing the effects of unstructured noise in the data and real cross-subject variability, not from alignment improvements. Notably, other fields concerned with image analysis generally strive to reduce rather than increase blurring of their images (e.g., astronomy uses space telescopes and advanced adaptive optics to counteract the blurring effects of the earth’s atmosphere).

Additionally, the use of statistical thresholds (which are less reproducible than effect size maps and spatial gradients (13)), and the tendency to summarize activation clusters as single 3D coordinates, induce further spatial localization uncertainty in the traditional volume-based literature. Taken together, these limitations make accurate comparisons between the traditional neuroimaging literature and the HCP’s multi-modal parcellation very challenging indeed. Our findings are also unlikely to be the result of circularity (i.e., that MSMAll was intrinsically favored because it had been used in generating the HCP-MMP1.0 parcellation used here to evaluate different acquisition and analysis methods) for the reasons discussed in Supplemental Discussion Section D1. Briefly, we used a spatially agnostic classifier, analyzed an independent dataset, and replicated the findings with a different version of the registration algorithm that produces a non-identical solution. Indeed any circularity concerns would be limited to the improvements of MSMAll over and above MSMSulc, which accounts for only ~25% of the 3-fold improvement of MSMAll over FNIRT with 8mm smoothing (~50% is related to not doing the smoothing and another ~25% to surface instead of volume alignment). MSMAll has previously been shown to provide major improvements in independent task fMRI data that was not used in the registration (20, 21) over MSMSulc. Thus, we believe any circularity effect is small if it exists at all, and even in a worst case it would not affect our main conclusions.

Our analysis focused on FSL’s FNIRT algorithm and alignment to the MNI152 FNIRT non-linear atlas template because that is how the HCP data were aligned volumetrically. Numerous other algorithms for volumetric alignment are available, and numerous template volumes are used as targets for registration to an atlas. However, we believe that no volume-based registration algorithm that enforces smoothly varying and anatomically plausible amounts of distortion will have better cortical alignment performance than the areal-feature-based surface registration algorithm shown here, even if areal features were used in the volume^3^. Volume-based registration of incompatible folding patterns and of incompatible relationships between areas and folds is a profoundly more challenging problem. While surface-based registration achieves a correspondence in 2D on a sphere, which allows spherical registration deformations to ignore incompatible or misleading folding patterns and therefore is unable to mix cortex with white matter or CSF, nonlinear volumetric registration is instead defined in terms of anatomical deformations, and must explicitly deal with fundamental folding issues like two gyri in some subjects where most have only one (see Supplemental Discussion Section D2 for further discussion of alternative volume-based registrations and the fundamental differences between surface and volume-based registration). On the other hand, even better correspondence across subjects than the current areal-feature-based surface registration may be attainable using improved algorithms within the MSM framework (20), increasing the amount of allowed distortion, or other topology-preserving approaches [e.g.,(39)]. Methods that do not enforce topology preservation such as individual subject parcellation (9, 40, 41) or hyperalignment (42) (36) can perform even better. Surface based studies, such as those that have used FreeSurfer alignment to the ‘fsaverage’ atlas, can profitably be compared to the HCP-MMP1.0 parcellation (see Supplemental Discussion Section D3). Indeed they are ~3/4ths of the way towards an HCP Style analysis (Figure 8). For legacy volume-based studies, it is now feasible to reanalyze such data using surface-based methods for accurate comparison with modern maps of the cerebral cortex (e.g. using tools such as CIFTIFY that are expressly designed for this purpose (https://github.com/edickie/ciftify)). Finally, it is worth remembering that while we believe the HCP’s multi-modal parcellation version 1.0 is the best available map of human cerebral cortical areas, we expect future refinements as more data become available and more investigators tackle the cortical parcellation problem using semi-automated, HCP-Style approaches (see Supplemental Discussion Section D4).

## Concluding Remarks

For decades, human neuroimaging studies have been dominated by an analysis paradigm consisting of volumetric alignment of cortical data coupled with unconstrained volumetric spatial smoothing. Unfortunately, this volumetric alignment is inaccurate for most of human neocortex, and results in statistically significant blobs whose precise relationship to cortical areas is uncertain. These blobs are then represented as 3D volumetric coordinates and assigned to Brodmann areas and/or coarse folding-related landmarks. Such results generally lack a close resemblance to the fine-grained mosaic of areas that populate the cortical sheet and generate the functional signals measured by these studies (9). Cortical surface-based approaches provide powerful alternatives that have gained momentum since their introduction two decades ago, especially for studies of visual cortex, e.g. (14), but widespread adoption of surface-based approaches has been slow [see (27); (13)], hampering progress in understanding cortical areas outside the visual system. Factors contributing to this unfortunate situation include the desire to replicate or compare with existing studies that used the traditional volume-based approach; the relative lack of “turn-key” tools for running a surface-based analysis [but see (19)]; the learning curve for adopting surface-based analysis methods; unawareness of the problems with traditional volume-based analysis; and uncertainty or even skepticism as to how much of a difference these methodological choices make.

The present study speaks mainly to the last two points, as we have used the HCP’s multi-modal parcellation to quantify the benefits of aligning data on surfaces instead of in volumes, using areal features instead of folds for this alignment, and minimizing spatial smoothing. These choices have a large impact on spatial localization. Moreover, analysis software and preprocessing pipelines for using ‘grayordinate-based’ analysis and visualization are now freely available, such as the HCP pipelines available on github (https://github.com/Washington-University/Pipelines), Connectome Workbench, growing support in FSL, and tools such as CIFTIFY for processing legacy data (https://github.com/edickie/ciftify). FreeSurfer has also provided its own surface-based analysis streams for nearly two decades (Fischl et al 1999). We hope that these approaches will receive increasing support and adoption in other tools such as AFNI (SUMA), BrainVoyager, and SPM (CAT).

Rather than continuing with “business as usual” using the traditional volume-based approach, we encourage neuroimaging investigators to re-evaluate the methods that they are using. An over-reliance on measures of statistical significance to assess scientific validity of brain imaging studies and an under-appreciation of neuroanatomical fundamentals has contributed to recent controversies in neuroimaging [e.g. (35)]. The issues considered in this and preceding papers [see (13)] are arguably even more problematic, given the vast resources poured into tens of thousands of neuroimaging studies that have yielded blurry results that would require re-analysis to enable accurate comparisons with the sharper picture that is emerging of the functional and structural organization of the cerebral cortex. The future impact of these legacy studies is significantly limited by the difficulty in accurately comparing them to modern surface-based maps of the human cerebral cortex.

Widespread adoption of approaches to data acquisition, analysis, and visualization that preserve high resolution and enable precise spatial localization is vital for accurately relating brain structure, function, and connectivity to well defined neuroanatomy. Progress in this direction will benefit from recognition and consideration of these issues by investigators, reviewers, journal editors, and funding agencies alike. Accurate relation of brain mapping results to brain areas can accelerate progress in understanding how the brain works in health and disease.

## Acknowledgements

We thank Alan Anticevic, Michael Harms, Erin Dickie, Babatunde Adeyemo, Takuya Hayashi, Bruce Fischl, Steve Smith, and Tom Nichols for comments and suggestions, along with three peer reviewers: Alex Cohen, James Haxby, and Martin Sereno. Supported by grants NIH F30 MH097312 (MFG), RO1 MH-60974 (DCVE), and 3R24 MH108315 (DCVE). Data were provided by the Human Connectome Project, WU-Minn Consortium (Principal Investigators: David Van Essen and Kamil Ugurbil; 1U54MH091657) funded by the 16 NIH Institutes and Centers that support the NIH Blueprint for Neuroscience Research; and by the McDonnell Center for Systems Neuroscience at Washington University.

## Footnotes

1 The only spatial constraint was that the classifiers were applied within searchlights that were large (30 mm in all directions from the group areal definition) relative to the size and variability of cortical area positions in areal-feature-based aligned data. Parcellation regularization steps that occurred after areal classification (dilation, remove islands, etc) used only local distance and vertex neighbor information.

2 Parcel-based averaging is appropriate for many analyses, but of course not for all. For example, studies of fine-scale cortical topography (e.g., 36-38) depend on preservation of small spatial details and are best analyzed using minimally smoothed data. However, neurobiological interpretation of topographic gradients can benefit from relating the gradients to nearby cortical areas.

3 That said, we feel volumetric registration of subcortical structures remains important (for example, the HCP uses the FNIRT nonlinear registration algorithm for subcortical alignment). Additionally the use of fiber orientation information may improve white matter fiber-tract alignment in the same way that areal-features improve cortical areal alignment (43-44). Additionally, the use of gentler volume registration algorithms optimized maximize functional alignment would be beneficial (see Supplementary Figures 11 and 12) together with including surface-based constraints that may also help resolve some ambiguities in the volume (45).

